# Anatomy of the Heart with the Highest Heart Rate

**DOI:** 10.1101/2021.10.11.463871

**Authors:** Yun Hee Chang, Boris I. Sheftel, Bjarke Jensen

**Affiliations:** Department of Medical Biology, University of Amsterdam, Amsterdam, Cardiovascular Sciences, Amsterdam UMC, Meibergdreef 15, 1105AZ, Amsterdam, The Netherlands; A.N. Severtsov Institute of Ecology and Evolution RAS (Russian Academy of Sciences). Leninsky prospect 33, Moscow, 119071, Russian Federation

**Keywords:** evolution, development, cardiac conduction system, trabeculation, pulmonary veins

## Abstract

Shrews occupy the lower extreme of the seven orders of magnitude mammals range in size. Their hearts are large relative to body weight and heart rate can exceed a thousand beats a minute. To investigate whether cardiac traits that are typical mammalian scale to these extremes, we assessed the heart of three species of shrew (genus *Sorex*) following the sequential segmental analysis developed for human hearts. Using micro-computed tomography we describe the overall structure and find, in agreement with previous studies, a large and elongate ventricle. The atrial and ventricular septums and the atrioventricular and arterial valves are typically mammalian. The ventricular walls comprise mostly compact myocardium and especially the right ventricle has few trabeculations on the luminal side. A developmental process of compaction is thought to reduce trabeculations in mammals, but in embryonic shrews the volume of trabeculations increase for every gestational stage, only slower than the compact volume. By expression of Hcn4, we identify a sinus node and an atrioventricular conduction axis which is continuous with the ventricular septal crest. Outstanding traits include pulmonary venous sleeve myocardium that reaches farther into the lungs than in any other mammals. Typical proportions of coronary arteries-to-aorta do not scale and the shrew coronary arteries are proportionally enormous, presumably to avoid the high resistance to blood flow of narrow vessels. In conclusion, most cardiac traits do scale to the miniscule shrews. The shrew heart, nevertheless, stands out by its relative size, elongation, proportionally large coronary vessels, and extent of pulmonary venous myocardium.

## INTRODUCTION

Shrews are eutherian insectivore mammals that can be tiny. Adult Eurasian least shrew (*Sorex minutissimus*) and Eurasian pygmy shrews (*Sorex minutus*), which we study in this report, only weigh a few gram. They therefore occupy the lower extreme of the seven orders of magnitude that mammals range in size. Because not much is known of their cardiac anatomy (Vornanen, 1989, Rowlatt, 1990), it is not clear whether typical traits of mammal hearts scale to such miniscule sizes. Valves and chamber wall thicknesses would be predicted to scale linearly to cavity size according to the law of Laplace (Seymour and Blaylock, 2000, Jensen 2021) and typical proportions of valves, walls, and cavities might well scale to shrews. In contrast, resistance to blood flow is inversely related to vessel diameter raised to the power of four according to the Hagen-Poiseuille equation and very small arteries and veins in the shrew may impose high resistance.

Blood perfusion constantly must be critical in shrews since with the tiny body size also comes with the highest rates of mass specific metabolism among mammals which has to be supported by the greatest mass specific cardiac outputs, or volume of blood per gram tissue pumped per minute (Morrison et al., 1959, Jurgens et al., 1996). Whereas in human it takes one minute to circulate the entire blood volume, in a shrew it takes a few seconds (Schmidt-Nielsen, 1984). A key component of the prodigious circulation is the incredibly fast heart rate, which may exceed a thousand beats a minute in the smaller shrew species (Morrison et al., 1959, Nagel, 1986, Vornanen, 1992, Jurgens et al., 1996). In many mammals, a well-developed cardiac conduction system initiates in its sinus node the cardiac impulse that, after a slow propagation through the atrioventricular node, rapidly spreads throughout the ventricles via the His bundle, bundle branches and Purkinje fibers (Davies, 1942, Dobrzynski et al., 2013). It is not known whether shrews have a typical conduction system, or an excessive one which could be expected given the extreme heart rates. An additional component in achieving great cardiac output is likely a relatively large stroke volume because heart mass is substantially greater in shrews than in most other mammals (Pucek, 1965, Bartels et al., 1979, Vornanen, 1989).

There is no consensus on how to analyze the gross anatomy of a mammal heart while it is a common approach to describe structures in the same order as blood flows through the heart (Kareinen et al., 2020, Marais and Crole). One similarly ordered and broadly used approach for human hearts is the sequential segmental analysis which is sufficiently versatile to be applicable to crocodile hearts (Cook et al., 2017). Here we followed this manner of analysis. Our specimens were caught in the wild, in fall traps in Siberia, Russia. By coincidence, a few of the trapped shrews were pregnant females. From the embryos of these, we could study the development of the ventricular walls. Early cardiogenesis has been described for the house shrew and demonstrates the presence of highly trabeculated ventricles (Yasui, 1992, Yasui, 1993), while anatomical studies on adult animals of the closely related moles of the genus *Talpa* suggest the adult ventricle may only have few trabeculations (Rowlatt, 1990). Gestational changes to ventricular trabeculations have attracted much attention in the context of so-called ‘noncompaction’ cardiomyopathy (Chin et al., 1990, Del Monte-Nieto et al., 2018, D’Silva and Jensen, 2020). Focus has been on a process of ‘compaction’, whereby trabeculated muscle on the luminal side of the ventricle is added to the compact wall (Sedmera et al., 2000). There is very little quantitative evidence, however, for such a process (Rychterova, 1971, Faber et al., 2021b). Instead, there is much stronger quantitative support for that the trabecular and compact layers can grow at different rates and when they do it changes the proportion of trabecular-to-compact myocardium (Blausen et al., 1990, Faber et al., 2021a, Faber et al., 2021c). If the shrew ventricles undergo pronounced gestational changes to the trabeculated layer, they may be a good test case for the explanatory power of compaction versus differential growth rates.

The primary research question of this study is whether typically traits of mammal hearts scale to the miniscule size of shrews. While studying the shrews, we had the opportunity to address the secondary questions of what is the state of their cardiac conduction system given the high heart rates and how do their ventricular walls develop.

## MATERIAL AND METHODS

### Animals

All animals were collected from near Yenisei Ecological Station of A.N. Severtsov Institute of Ecology and Evolution RAS, Turukhansk district of Krasnoyarsk region (N 62°17’, E 89°02’). The collection of animals and the manner of trapping them complied with the guidelines of the A.N. Severtsov Institute of Ecology and Evolution RAS. Briefly, animals were collected from water-filled fall traps, deskinned, and fixed in either 10% or 20% formalin for one day and then kept in 70% ethanol until further use. Of the Eurasian pygmy shrew (*Sorex minutus*), we used six heart-lung preparations for the description of the formed heart and two embryos from each of three pregnant females for the description of developing hearts (Table 1). In addition, we used heart-lung preparations of formed animals of two Eurasian least shrew (*Sorex minutissimus*) and two taiga shrew (*Sorex isodon*) (Table 1).

**Table 1.**
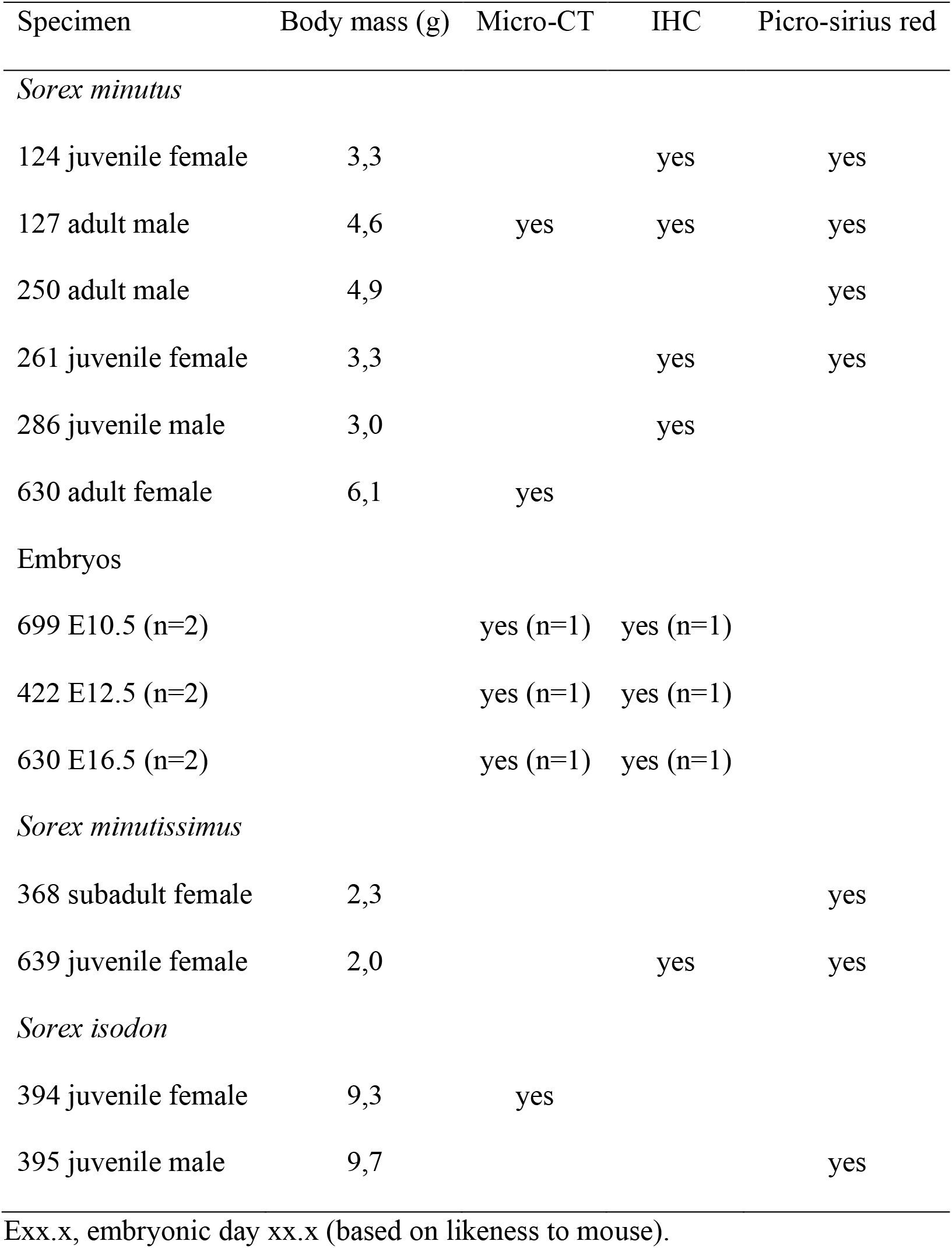
Overview of the used specimens.

### Micro-computed tomography

The three hearts that were investigated with micro-CT, two *Sorex minutus* and one *Sorex isodon*, were first stained for two days with Lugol’s solution (1.75g I_2_ and 2.50g KI, both from Fischer Scientific, dissolved in 100ml deionized water). The volume of Lugol’s solution was at least 10 times that of the tissue and the solution was kept in the dark during the staining. The tissue was then immobilized by imbedding in an agar solution (1.3g per 100ml water). Subsequently they were scanned at isotropic 10 μm resolution using a Bruker, Skyscan 1272. All shown images of micro-CT are of specimen 127 which had the best tissue-lumen contrast of the two *Sorex minutus* specimens.

### Sectioning and Histology

The hearts that were investigated with histology were embedded in paraplast and cut in 10 or 12 μm sections in either the transverse or frontal plane (four-chamber view). Staining was with saturated picro-sirius red in which muscle becomes orange and collagen becomes red following 2 min differentiation in 0.01M HCl. Imaging of the stained slides was done with a Leica DM5000 light microscope.

### Immunohistochemistry

With fluorescent immunohistochemistry, we detected smooth muscle actin (SMA), cardiac troponin I (cTnI) and hyperpolarization activated cyclic nucleotide gated potassium channel 4 (Hcn4) as well as nuclei using DAPI (Sigma-Aldrich, dilution 1:1000, D9542). For SMA we used the primary antibody RRID:AB_476701 (Sigma-Aldrich, dilution 1:400) which was visualized by a fluorescently labeled secondary donkey anti-mouse antibody coupled to Alexa 555 (Thermo Fisher Scientific, dilution 1:250, RRID:AB_2536180). For cTnI we used the primary antibody RRID:AB_154084 (Hytest, dilution 1:300) which was visualized by a fluorescently labeled secondary donkey anti-goat antibody coupled to Alexa 488 (Thermo Fisher Scientific, dilution 1:250, RRID:AB_2762838). For Hcn4 we used the primary antibody RRID:AB_2120042 (Millipore, dilution 1:200) which was visualized by a fluorescently labeled secondary donkey anti-rabbit antibody coupled to Alexa 647 (Thermo Fisher Scientific, dilution 1:250, RRID:AB_2536183). In addition, for identification of the sinus node we tested Isl1 (RRID:AB_2126323) and Shox2 (RRID:AB_945451) but we never detected positive nuclei. We did not seek to clarify whether the absence of signal was due to absence of the protein or deterioration of the epitopes due to suboptimal tissue fixation and preservation. Slides were viewed and photographed with a Leica DM6000B fluorescent microscope. For estimations of the volume of trabecular and compact myocardium in embryos, we stained for cTnI as above on one section of 10μm thickness per 100μm (9 sections for specimen 699; 8 sections for specimen 422; 14 sections for specimen 630).

### Analyses and statistics

We imported to Amira (version 2020.2, ThermoFisher Scientific) the micro-CT image series (two specimens of formed heart and lungs, one embryo of each the three gestational ages) and the images used for estimation of ventricular trabecular and compact tissue volume (one embryo of each the three gestational ages). Concerning *Sorex minutissimus*, from the histological section series of the entire heart and lung preparation we imported to Amira every 25^th^ section, in total 13 sections for specimen 368 and 14 sections for specimen 639. Structures of interest were then labelled and their volume was derived using the “Materials Statistics” tool. To measure cross-sectional areas of vessels we first used the Slice module to create an image plane perpendicular to the labelled vessel. That plane was then converted to a 8-bit label image. Using the module Label Analysis, the vessel area was measured. To measure distances of structures of the lungs we imported images to ImageJ (version IJ 1.46r) and used the straight line selection tool. We used a one-way ANOVA to test for differences in distance.

## RESULTS

### Heart position and orientation

The body mass differed between the three investigated species (Table 1) and so did the size of the heart. The hearts were proportionally large, their tissue volume comprised more than 0.7% of fresh body mass (Table 2) and this percentage would likely have been greater still if fresh hearts had been measured (Vornanen, 1989). Besides the difference in absolute size of the heart between the species, there were no gross morphological differences between the hearts. The position of the shrew heart is much like in human (Fig. 1). Within the thoracic cavity, the heart is mostly on the left side, and immediately caudal to the heart is the diaphragm. The ventricles in particular are quite elongate and the rather pointy apex is made up of the apex of the left ventricle (LV) (Fig. 1A-D). There is no interventricular sulcus and the border between the left and right ventricle is subtle. The left atrium (LA) is the most cranial and dorsal chamber, the right atrium (RA) is the right-most chamber, and the right ventricle (RV) rests on the diaphragm and is the most caudal and ventral chamber (Fig. 1C). The LV is longer and narrower than the RV, with the length/breadth index of the left and right ventricle being 1.5 and 1.3, respectively (Fig. 1D-E). Three caval veins connect to the RA (Fig. 1F-G). Two pulmonary veins connect to the LA, of which the right pulmonary vein is wider as it is the confluence of the veins from each of the four lobes of the right lung (Fig. 1F,H).

**Figure 1.**
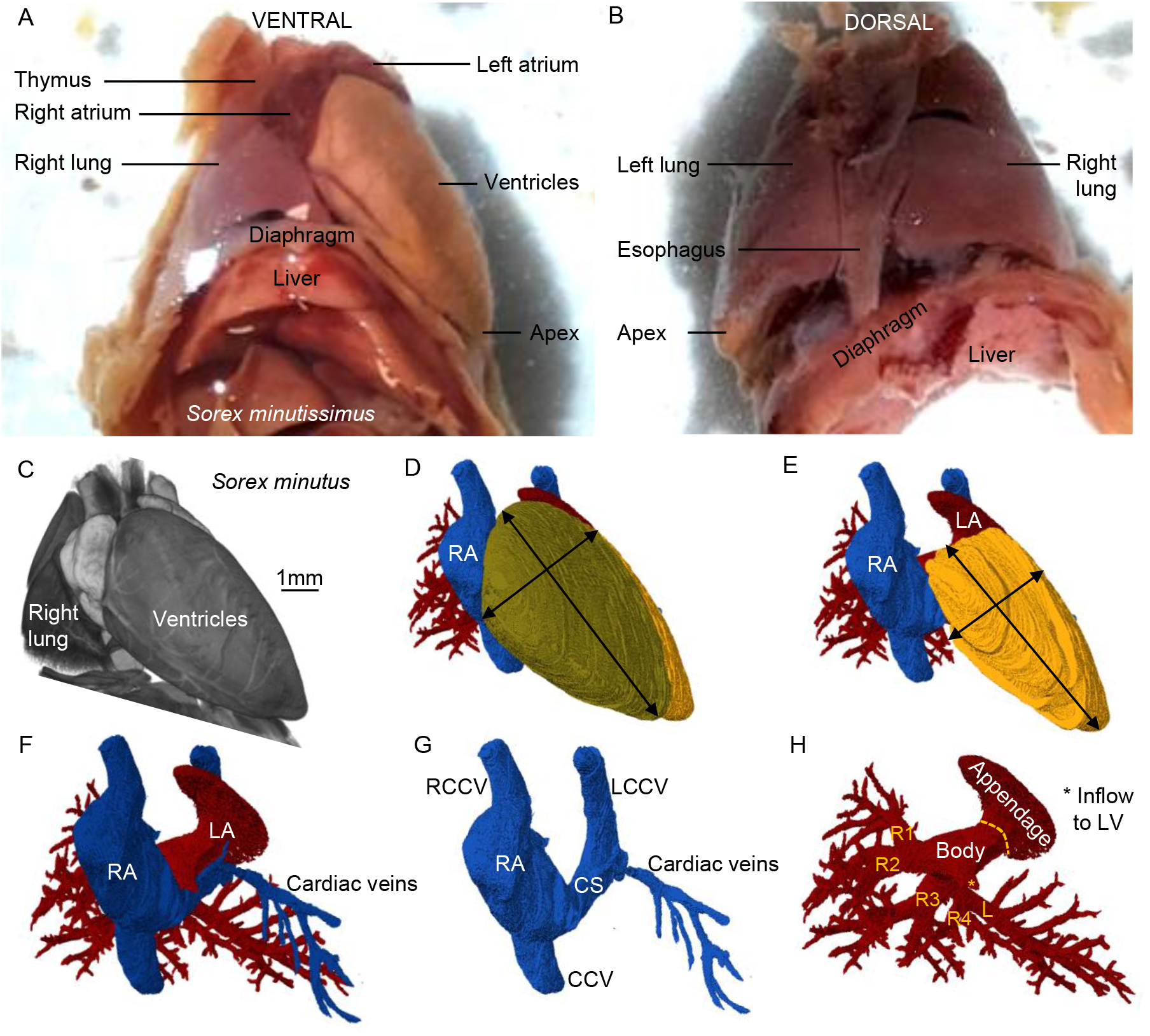
Topology of the shrew heart. **A-B**. Ventral (A) and dorsal (B) view of the heart of *Sorex minutissimus* (specimen 637). Notice the caudal-left position of the apex and the great elongation of the ventricle. **C**. Volume rendering of micro-CT of *Sorex minutus* (specimen 127), showing the heart in approximately its attitudinally correct position when viewed ventrally. **D**. The right atrium (RA) and right ventricle (RV) are the most ventral chambers. **E**. The left ventricle is very elongate. **F-G**. Three caval veins connect to the RA. **H**. The left atrium (LA) has a large appendage. It receives one stem into which drains veins from the first two lobes of the right lung (R1-R2) and a second stem into which drains veins from the last two lobes of the right lung (R3-R4) and the left lung (L). CCV, caudal caval vein; CS, coronary sinus; L(R)CCV, left (right) cranial caval vein.

**Table 2.** The volume of the heart and the pulmonary venous myocardium.

| Specimen | Body mass (g) | Cardiac volume (mm<sup>3</sup>) | Cardiac index (%) | PVM volume (mm<sup>3</sup>) | PVM/cardiac volume (%) |
| --- | --- | --- | --- | --- | --- |
| *Sorex minutus* |  |  |  |  |  |
| 286 | 3.0 | 23.34 | 0.78 | 0.488 | 2.09 |
| 261 | 3.9 | 28.42 | 0.73 | 0.360 | 1.27 |
| 127 | 4.0 | 35.70 | 0.78 | 1.163 | 3.26 |
| *Sorex minutissimus* |  |  |  |  |  |
| 368 | 2.3 | 16.02 | 0.70 |  |  |
| 639 | 2.0 | 15.92 | 0.80 |  |  |

PVM, pulmonary venous myocardium.

### The right side of the heart

#### Caval veins

Three caval veins connect to the RA, namely the caudal caval vein (CCV), right cranial caval vein (RCCV), and the left cranial caval vein (LCCV) (Fig. 1F-G). The part of the LCCV that is most proximal to the RA could be considered the equivalent of the coronary sinus of the human heart as it receives the great cardiac veins, it lies in the left atrioventricular (AV) groove and opens into the vestibule of the RA (Fig. 1G). There is no bridging vein between the RCCV and the LCCV, while such vein can be found in human with persistent left superior caval vein (Kula et al., 2011). The cross-sectional area of the lumens of the caval veins is large and comprises more than 50% of the total cross-sectional area of the lumens of all great veins and arteries (Table 3). All three caval veins have myocardium extending to the pericardial reflection (Fig. 2A-C), whereas in human there is hardly any myocardium in the inferior caval vein (Noheria et al., 2013). The distal-most parts of the cranial veins where often not included in the excised preparations, but when they were we found venous valves cranial to the myocardial-venous boundary. Also, at the myocardial-venous boundary of the caudal caval vein we found a valve in all three species (Fig. 2D-E).

**Figure 2.**
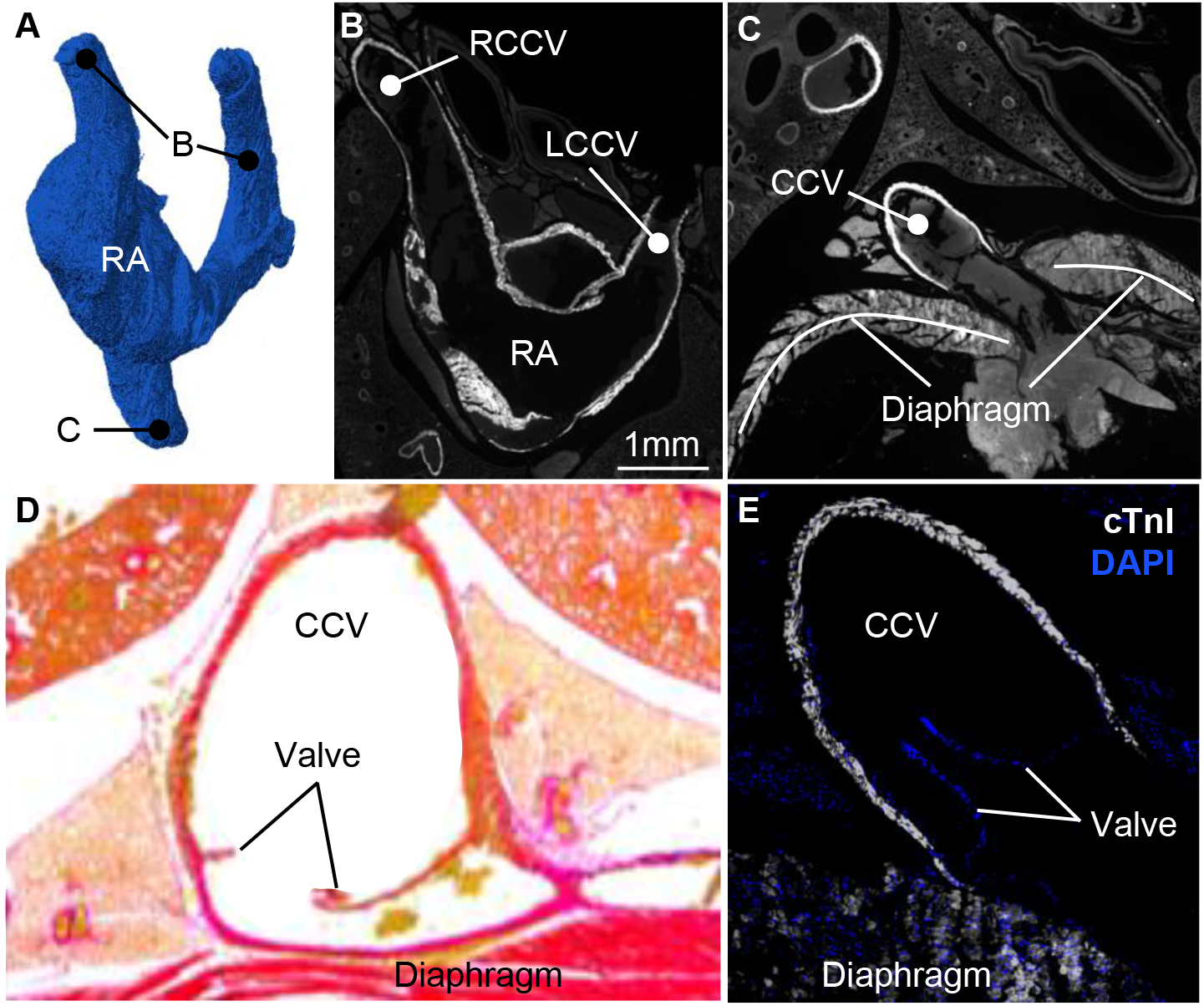
Extensive myocardial sleeves of the caval veins. **A**. Lumen cast of the right atrium (RA) and caval veins. Label B points to the approximate cranial-most extent of the myocardial sleeves which is documented in image **B**. Label C points to the caudal-most extent of the myocardial sleeves which is documented in image **C. B**. Detection of cTnI in the left and right cranial caval vein (LCCV and RCCV respectively). **C**. Detection of cTnI in the caudal caval vein (CCV). **D**. Venous valve at the myocardial-venous boundary in *S. minutissimus* (for the sake of presentation, blood inside the vein has been painted over with white). **E**. Venous valve at the myocardial-venous boundary in *S. minutus*.

**Table 3.**
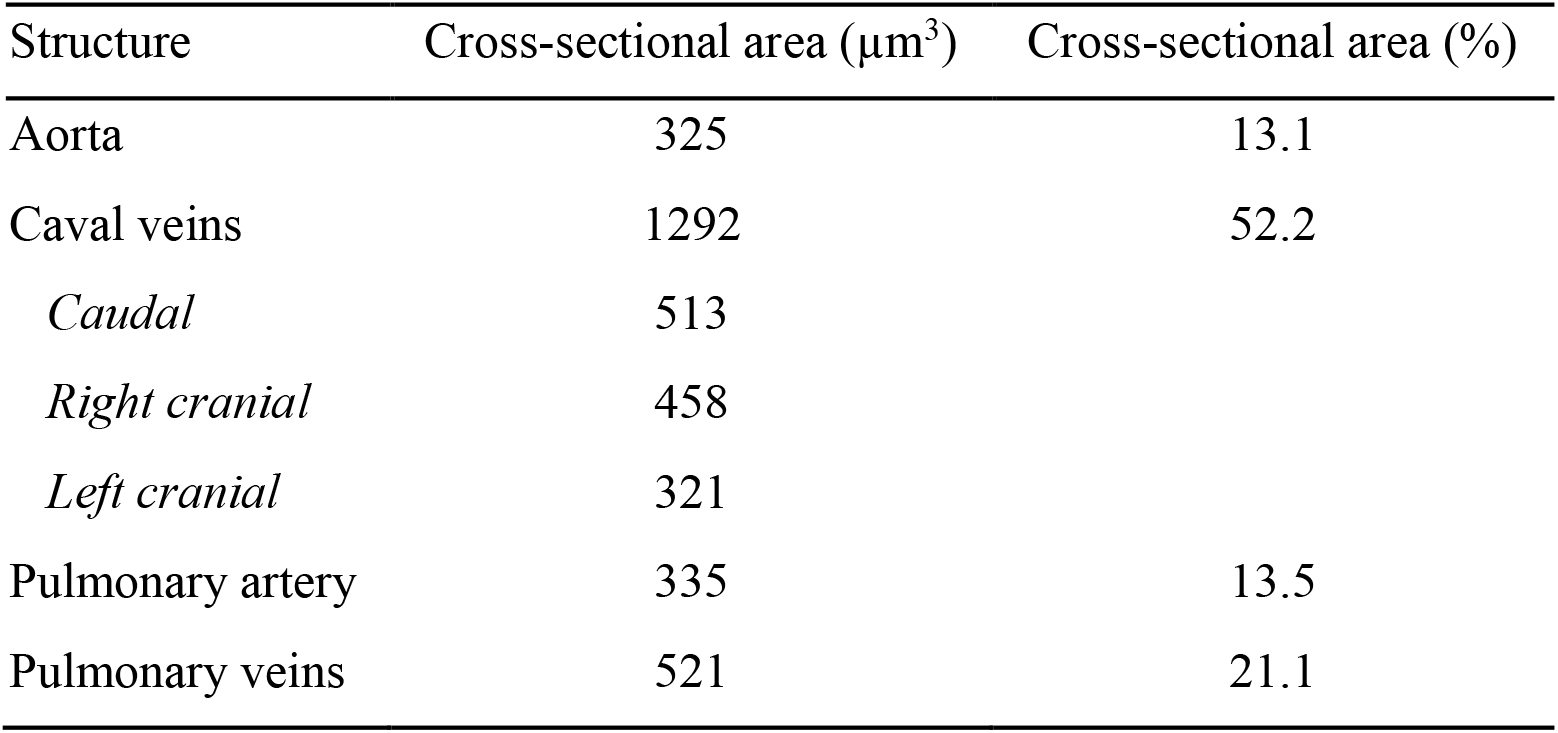
The cross-sectional area of the great vessels.

| Structure | Cross-sectional area (μm<sup>3</sup>) | Cross-sectional area (%) |
| --- | --- | --- |
| Aorta | 325 | 13.1 |
| Caval veins | 1292 | 52.2 |
| *Caudal* | 513 |  |
| *Right cranial* | 458 |  |
| *Left cranial* | 321 |  |
| Pulmonary artery | 335 | 13.5 |
| Pulmonary veins | 521 | 21.1 |

#### The right atrium and atrial septum

A very prominent pillar-shaped muscular ridge, the crista terminalis is found in the dorsolateral wall of the RA, where it divides the atrium into a dorsal smooth portion, or body, and a lateral trabeculated portion, or appendage (Fig. 3A). The trabeculations of the body are thin compared to the LA (see below) and, when compared to the human RA, the trabeculations are configured more as a network than parallel pectinate muscles. Also, in contrast to human, the body is bigger than the appendage (Fig. 3B). The trabeculation extends to the area immediately around the right AV valve and atrial septum, except for the area of the vestibule. There is a well-developed left leaflet of the sinuatrial valve, whereas the right leaflet is much less prominent (Fig. 3C). The atrial septum comprise a thick secondary atrial septum composed of predominantly myocardium and a thin primary septum, or flap valve, composed of some myocardium besides endocardium and connective tissue (Fig. 3D). The cranial aspect of the secondary septum, which in human is essentially a fold (Anderson et al., 2014), in the shrew also takes the appearance of a folding-in of the atrial roof albeit the sulcus is traversed by bits of myocardium.

**Figure 3.**
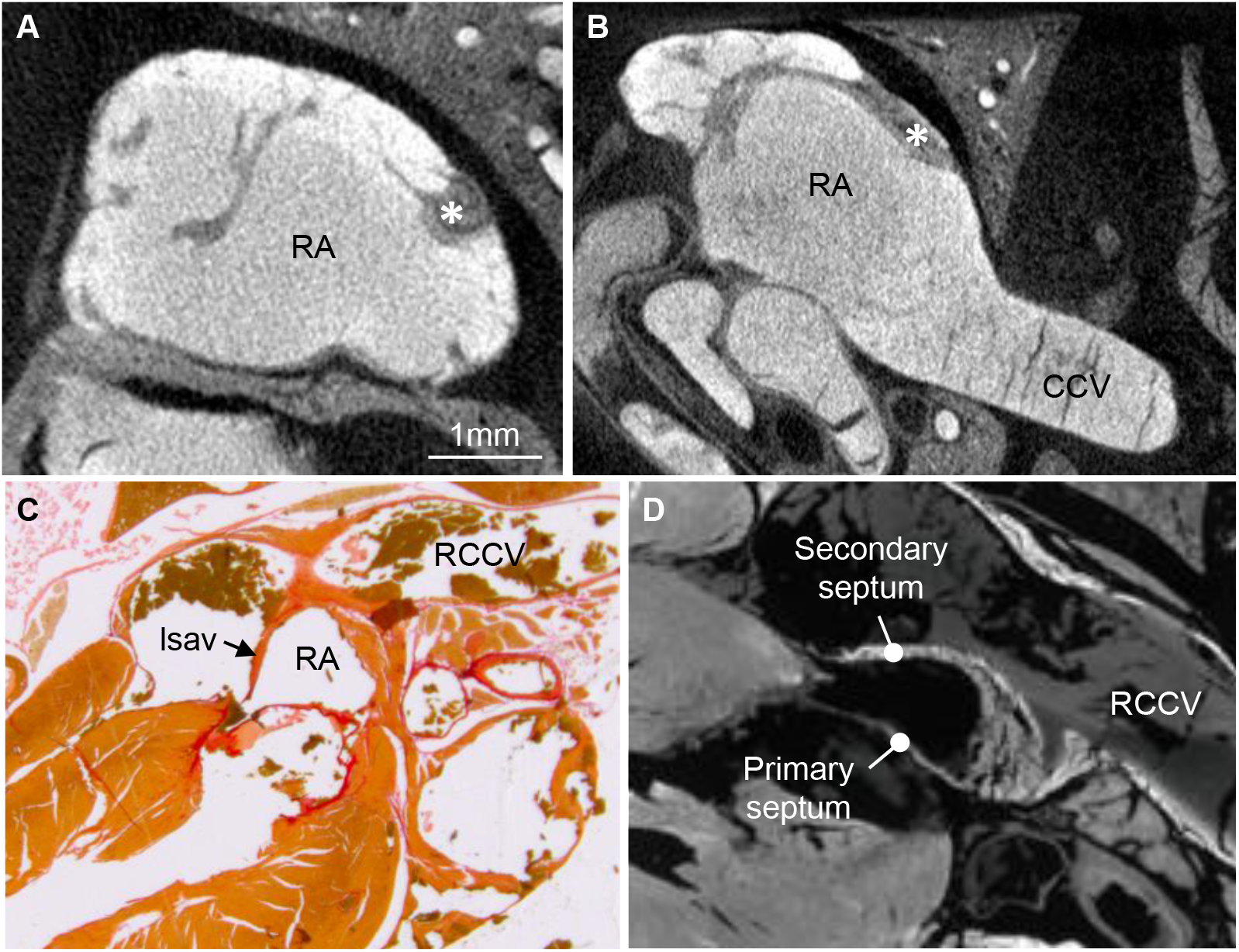
Anatomy of the right atrium. **A**. The wall of the right atrium (RA) comprise a meshwork of trabeculations, the greatest among which is crista terminalis (asterisk). **B**. There is a comparatively extensive smooth-walled part, or body. **C**. A well-developed left leaflet of the sinuatrial valve (lsav) extends from the junction of the right cranial caval vein (RCCV) to the base of the atrial septum. **D**. The atrial septum has a primary component that is thin and contains much non-myocardial tissue and a secondary component mostly of myocardium. CCV, caudal caval vein.

#### The right ventricle and pulmonary artery

The right ventricle looks to wrap around the septal part of the left ventricle (Fig. 4A) as is typical in mammals. The atrioventricular valve is membranous and of connective tissue (Fig. 4B). It comprises a large septal leaflet and a large parietal leaflet (Fig. 4C). A prominent cleft subdivides the parietal leaflet in a smaller dorsal part and a large ventral part (Fig. 4C). The septal surface has a mostly smooth surface. Consequently, the appearance of the papillary muscles and chordae tendineae is very subtle (Fig. 4A). Most the luminal side of the ventricular wall in fact has a smooth, or a-trabecular, appearance including the outflow tract. At the transition from septum to wall, however, there are well-developed trabeculations (Fig. 4D). Also, a fairly prominent trabecula septo-marginalis was found in the typical mammalian position (ventro-apically) and it hosted a proportionally quite large coronary artery (Fig. 4E). There are three cusps to the pulmonary arterial valve. As in human, these hinge in ventricular myocardium that forms a myocardial turret around the base of the pulmonary artery (Fig. 4F).

**Figure 4.**
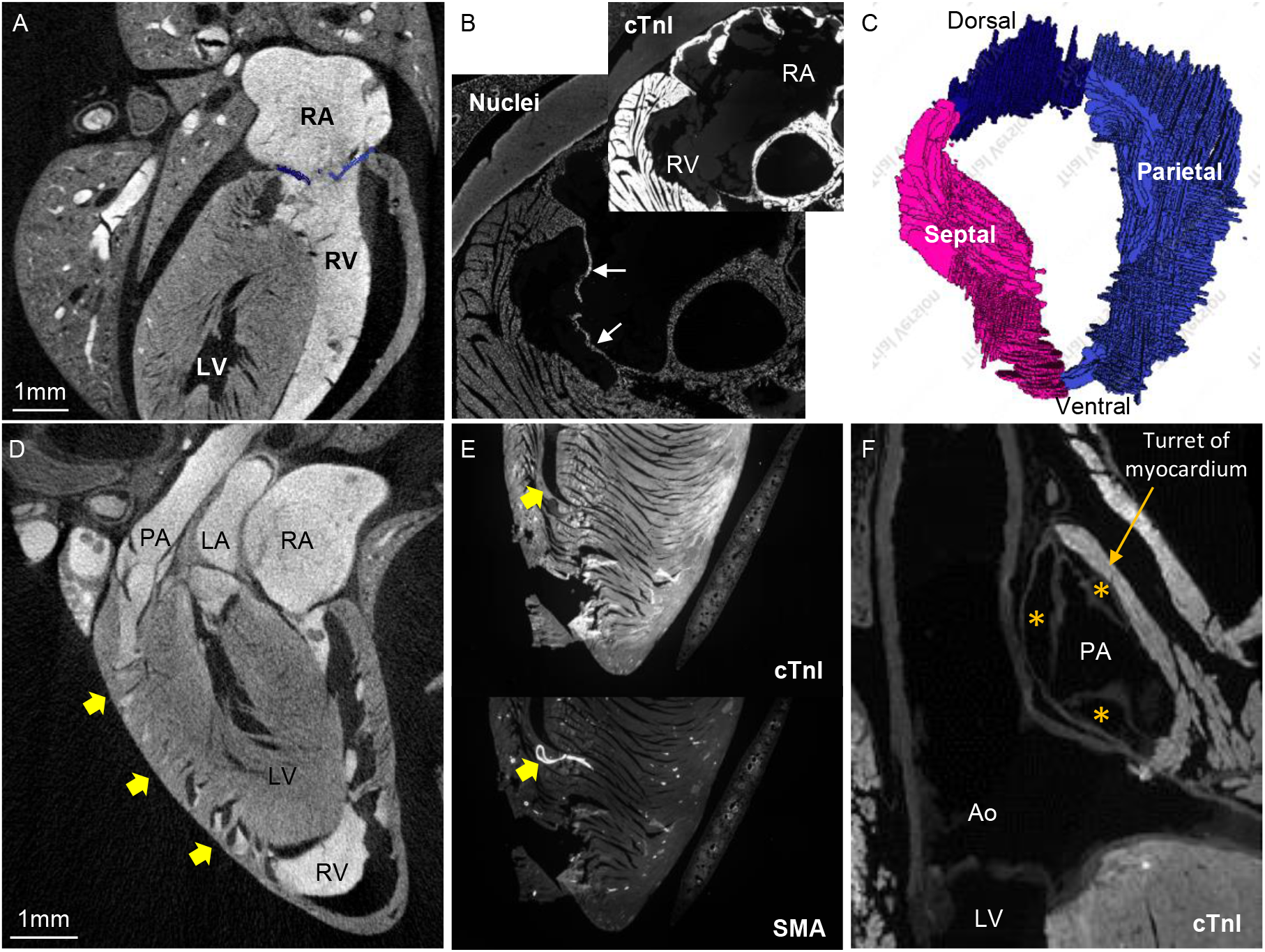
Anatomy of the right ventricle. **A**. The right ventricle (RV) wraps around the left ventricle (LV). Notice the smooth appearance of the septal surface and parietal wall, including an absence of prominent papillary muscles. **B**. The atrioventricular valve is membranous (white arrows) and without myocardium. **C**. Reconstruction of the atrioventricular valve, showing a prominent septal leaflet and a parietal leaflet that has a cleft such that it is divided into two parts. **D**. At the boundary between wall and septum are numerous trabeculations (arrows). **E**. Trabecula septo-marginalis and moderator band (arrow), within which there is a large coronary artery (its walls contain smooth muscle actin (SMA)). **F**. The valve of the pulmonary artery (PA) has three leaflets that hinge in a ‘turret’ of myocardium.

The main trunk of the pulmonary artery is short and it bifurcates into one branch to each lung (Fig. 5). Just as the right lung is bigger than the left, the right branch of the pulmonary artery is bigger than the left. The right branch lies between the ascending and descending aorta and splits into 4 branches, one branch for each of the four lobes of the right lung (Fig. 5A-B). The smaller solitary left pulmonary artery has an angle of approximately 75° to the main trunk and goes into the left lung which has one lobe only (Fig. 5C-D). The aorta and the main trunk of the pulmonary artery are conduits for the same cardiac output and they have similar cross-sectional areas (Table 3).

**Figure 5.**
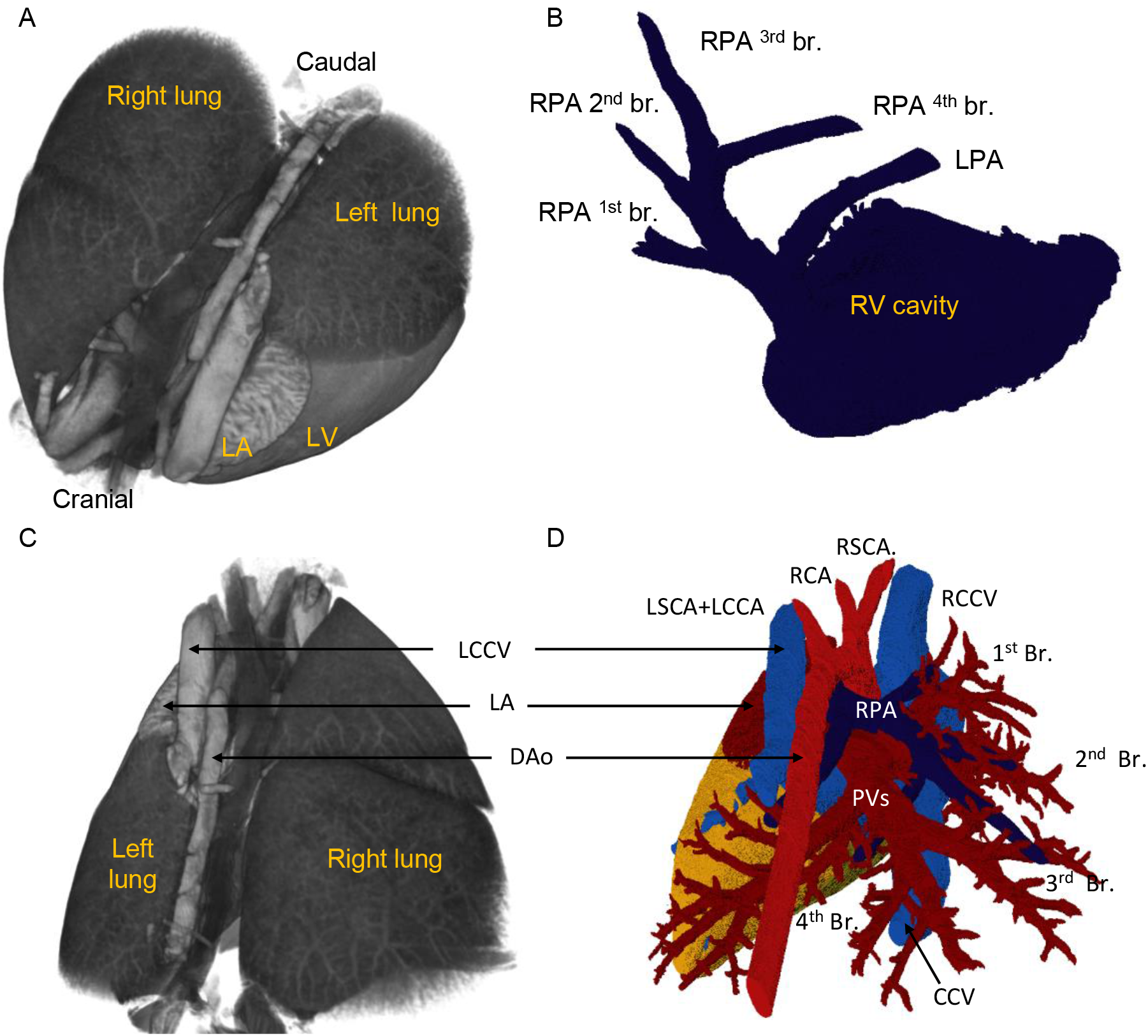
The pulmonary artery. **A**. Silhouette (blue) of the right ventricle and main branches of the pulmonary artery unto a volume rendering, based on micro-CT, of the heart and lungs. **B**. Virtual lumen cast of the cavity of the right ventricle (RV) and the pulmonary artery showing the four main branches to the right pulmonary artery (RPA 1^st^-4^th^) and the solitary left pulmonary artery (LPA). **C-D**. Dorsal view of the pulmonary circulation.

#### The ventricular septal structures

A small membranous septum was found below between the base of the aorta and the crest of the myocardial ventricular septum. The right atrioventricular valve hinges onto the membranous septum and thereby divides into an atrioventricular component (it separates the cavities of the right atrium and the left ventricle) and an interventricular component (Fig. 6A). A substantially offset in the hinge-line for the tricuspid valve relative to the mitral valve hinge-line was not seen. The atrioventricular membranous septum occupies the base of the gap between the non-coronary and right coronary cusps of the aortic valve (Fig. 6B-C).

**Figure 6.**
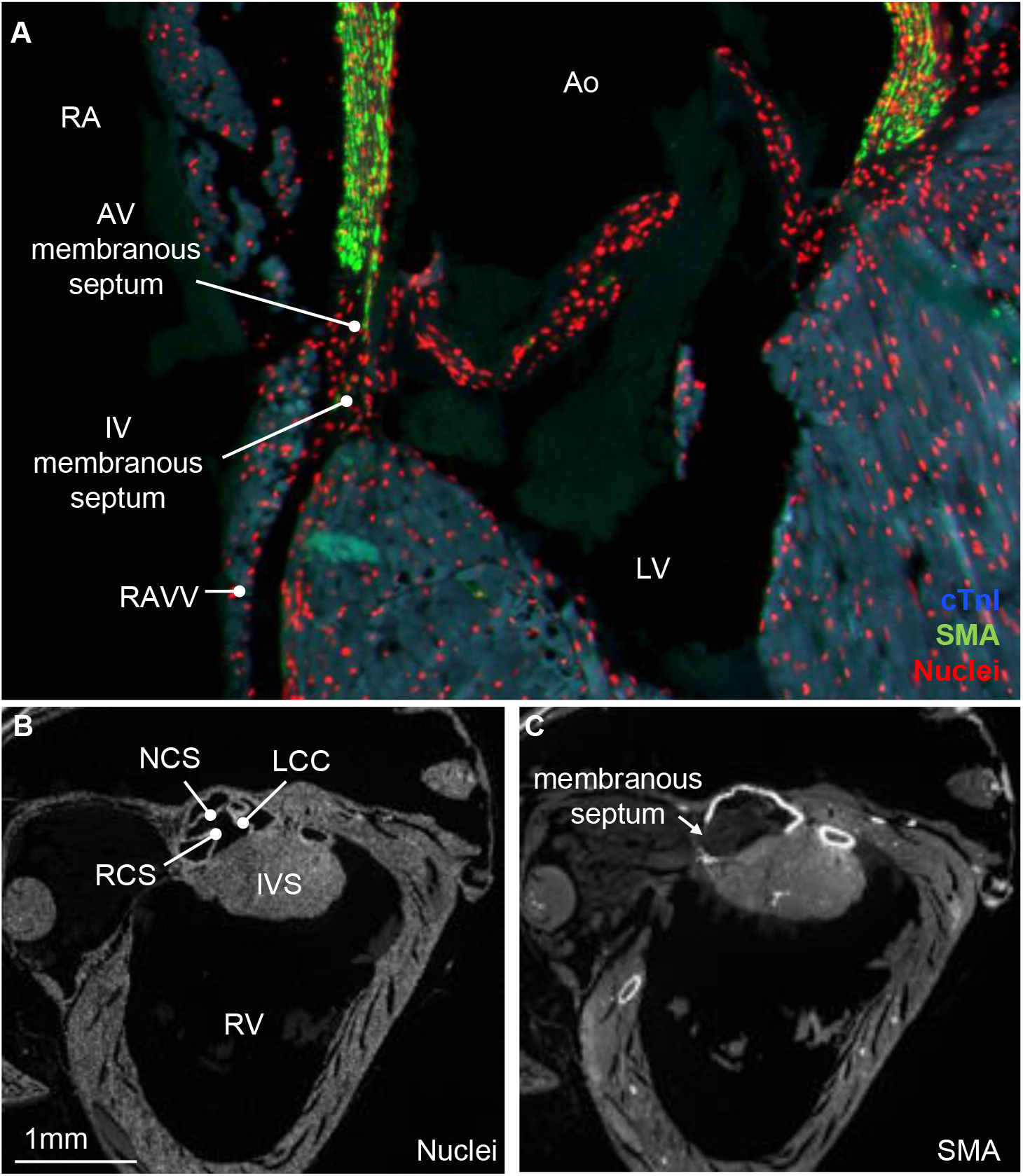
The membranous septum. **A**. The small membranous septum (cTnI negative) connects with the septal leaflet of the right atrioventricular valve (RAVV). Above this hinge line is the atrioventricular (AV) membranous septum and below it is the interventricular (IV) membranous septum. **B-C**. The atrioventricular membranous septum is below the base of the non-coronary (NCC) and right coronary cusps (RCC) of the aortic valve. Ao, aorta; LCC, left coronary cusp; LV, left ventricle; RV, right ventricle.

#### Pulmonary veins

The left-sided atrial body receives two very short stems of pulmonary veins, with the right stem coming from the first two lobes of the right lung and the left stem coming from the third and fourth lobe of the right lung together with the solitary vein of the left lung (Fig. 1F). The right stem passes between the entrance of the right cranial caval vein and caudal caval vein and left stem passes over the coronary sinus to reach the left atrium.

A remarkable feature of the shrew heart is the extent of the myocardial sleeves of the pulmonary veins (Fig. 7). In *S. minutissimus* and *S. minutus* these sleeves reach within 0.2 mm of the lung surface, even in the distal parts of the lungs (Fig. 7C-D). Measured as the distance to the lung surface, the pulmonary venous myocardium extends as far as the pulmonary arteries and the terminal parts of the alveolar ducts (Fig. 7C-D). Despite the great extent of the pulmonary venous myocardium, it only constitutes approximately 2% of total myocardial volume (Table 2).

**Figure 7.**
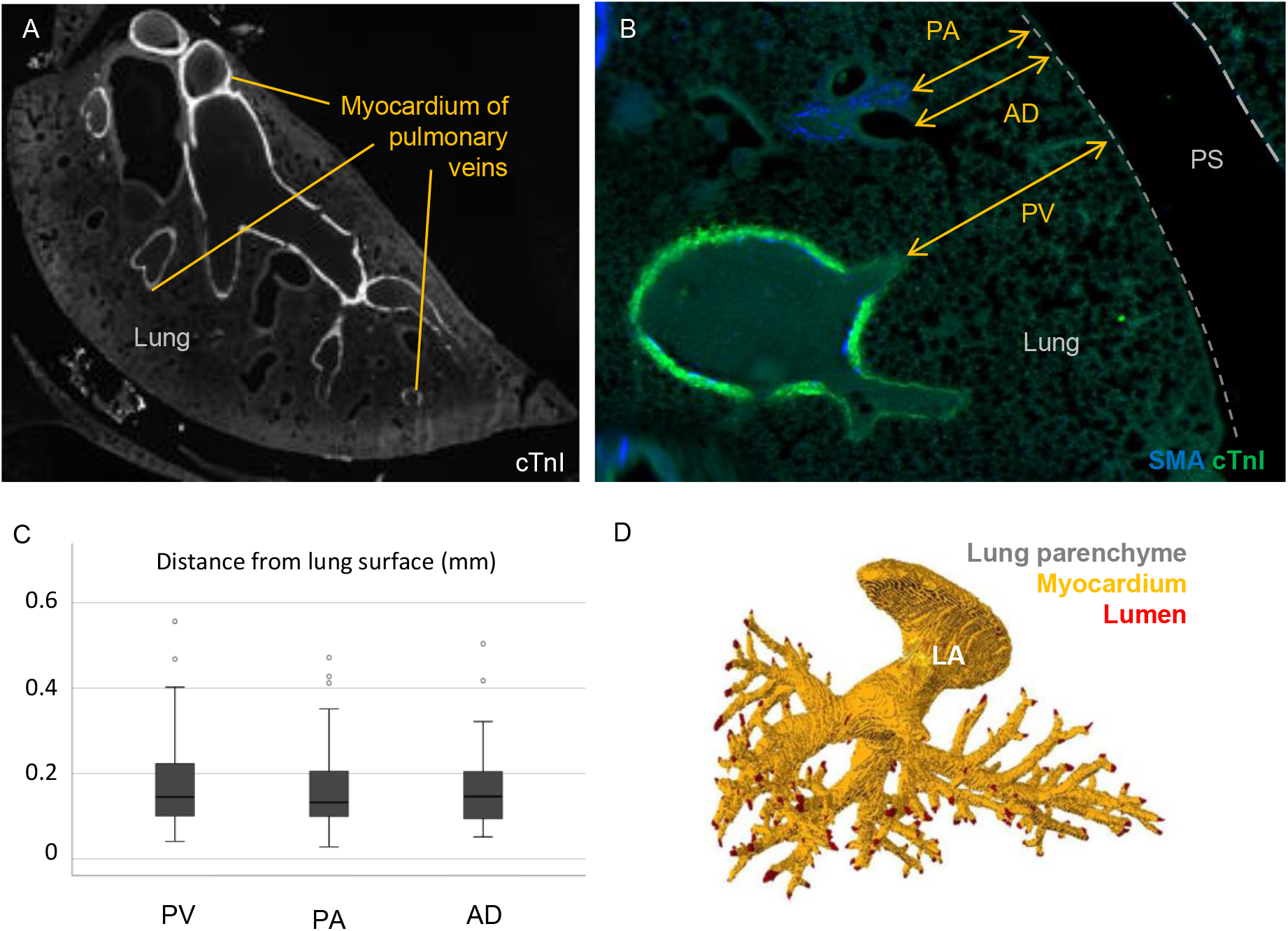
Extreme extent of pulmonary venous myocardial sleeves. **A**. Myocardium in the walls of the pulmonary veins reached the farthest parts of the lungs. **B-C**. Detailed view showing the pulmonary veins (PV), alveolar ducts (AD) and pulmonary arteries (PA) all extended to the proximity of the lung surface, and the distance of this proximity was not different on average (**C**, one-way ANOVA, p=0,573; 3 specimens, 4 or 5 sections per specimen, 53 measurements per structure). **D**. Reconstruction of the pulmonary venous lumen (red) and myocardial sleeves, illustrating that only the distal most parts of the pulmonary veins were without sleeves. PS, pleural space.

#### Left atrium

The left atrium is situated dorso-cranially. It has a venous component (body), a trabeculated component (appendage), and a vestibule (Fig. 8). The body has the two orifices of the pulmonary veins. The appendage is proportionally large when compared to the human setting. It is clog-shaped and its junction with the body is relatively narrow (Fig. 1F). The pectinate muscles are much more extensive in the left atrial appendage than the right atrial appendage (Fig. 8). Even though the left atrium lacks the muscular bundle like crista terminalis in the right atrium, the junction between the appendage and body is well defined by the coarse trabeculation in the appendage (Fig. 8).

**Figure 8.**
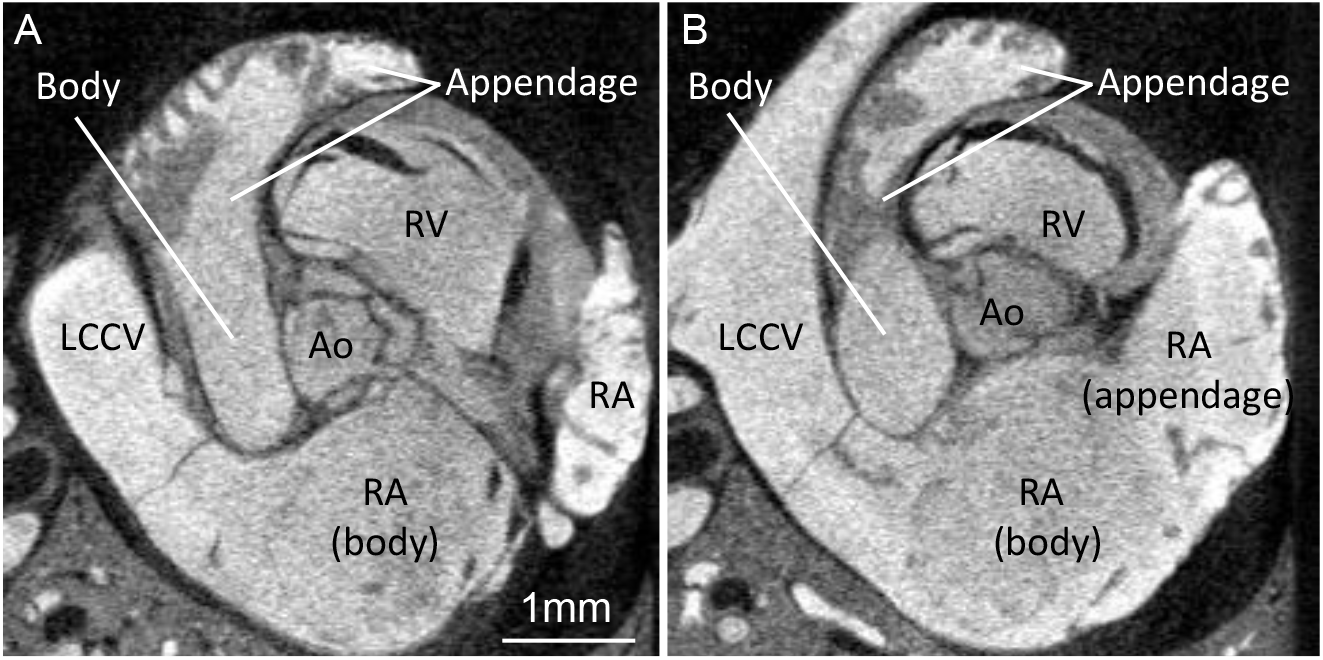
The left atrium. **A**. The left atrium has a smooth-walled body and a proportionally large trabeculated appendage. **B**. Trabeculations of the left atrium are more coarse than those of the right atrium (RA). Ao, aorta; LCCV, left cranial caval vein; RV, right ventricle.

#### Left ventricle and aorta

The left atrioventricular valve has two leaflets and two papillary muscles (Fig. 9). From the valve margins, chordae tendineae connect to the ventral papillary muscle which emerges from the ventricular septum and to the dorsal papillary muscle that emerges from the ventricular free wall (Fig. 9B-C). The papillary muscles and tension apparatus of the left ventricle are much more prominent than those of the right ventricle. With the exception of the septo-parietal trabeculation of the right ventricle, the trabeculations of the left ventricle are coarser than those of the right ventricle, and this setting therefore resembles that of pig but not that of human where the right ventricle has the coarser trabeculations (Crick et al., 1998). The left ventricular outflow tract is without trabeculations and the septal atrioventricular leaflet and fibrous continuity are also the part of the outflow tract (Figs. 6A, 9A).

**Figure 9.**
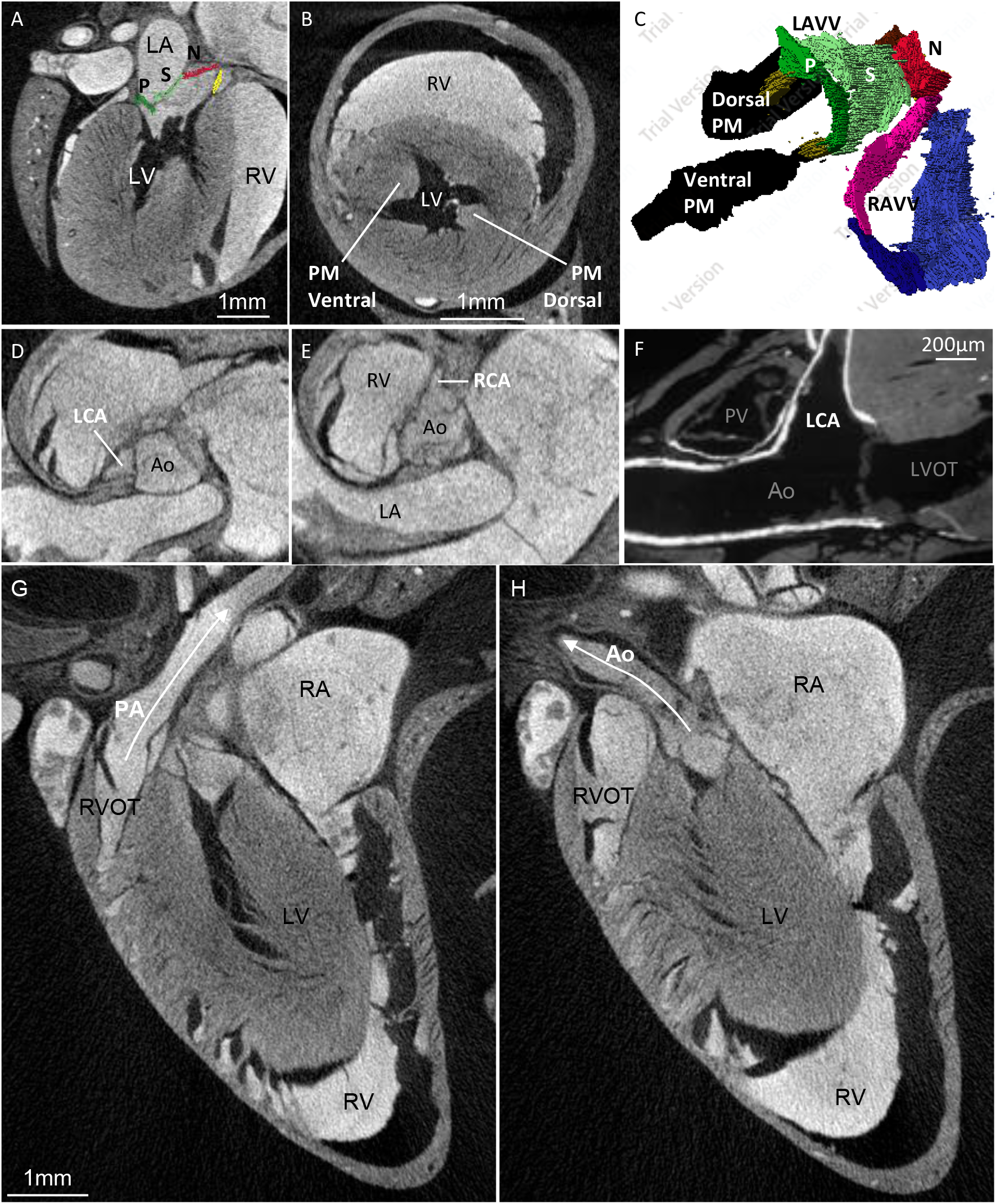
The left ventricle and aorta. **A**. The left atrioventricular valve has a parietal leaflet (P) and a septal leaflet (S) which is continuous with the aortic leaflet of the non-coronary sinus (N). **B**. There are two large papillary muscles (PM). **C**. Reconstruction of the atrioventricular valve and tension apparatus, the latter of which is much more developed from the left atrioventricular valve (LAVV) than the right atrioventricular valve (RAVV). **D-E**. Origins of the left (LCA) and right coronary artery stem (RCA) within the sinuses of the aortic valve. **F**. The LCA is proportionally very large, its diameter is almost a half of the diameter of the aorta (Ao) and the base of the pulmonary arterial valve (PV). **G-H**. The aorta and the pulmonary artery are oriented at very different angles as is typical of mammals. L(R)A, left (right) atrium; L(R)VOT, left (right) ventricular outflow tract; L(R)V, left (right) ventricle.

There are three cusps to the aortic valve and each of the two coronary arteries arise from its own aortic sinus like in human (Fig. 9D-E). The left coronary artery has a proportionally very large diameter, approximately 1:2 when compared to that of the aorta, and this ratio in human would be much closer to 1:10 (Fig. 9F). Its actual diameter is only approximately 200μm, however, which is the diameter of an arteriole. The right coronary artery is almost as large as the left. Both coronary arteries immediately become intramural in their course through the ventricular mass and their main stems are in the parietal walls rather than at the boundary of the left and right ventricle. Consequently, the ventricular surface does not have large coronary vessels and an interventricular sulcus that demarcate the left and right ventricle as in human and pig (Crick et al., 1998).

The aorta and the pulmonary trunk have a spiral relationship (Fig. 9G-H). The aortic arch crosses cranially to the bifurcation of the pulmonary arteries (Fig. 5D). The ascending aorta is located to the right of the trunk of the pulmonary artery. The aortic arch gives rise to only two branches leading cranially. The descending aorta lies between the esophagus and left cranial caval vein (Fig. 5D).

#### Development of ventricular trabeculation

The smooth appearance of the ventricular walls, suggested that extensive compaction had taken place during development, where compaction can be defined as a reduction of the trabecular layer as trabeculations are added to the compact wall (Faber et al., 2021b). To detect compaction, we investigated the gestational change to the volumes of the trabecular and compact layers of both ventricles, in hearts that by outward appearance resembled hearts of mouse from gestational ages E10.5, E12.5, and E16.5 (de Boer et al., 2012). The shrew heart grows rapidly (Fig. 10A), as seen in developing human, mouse and chicken as well (Faber et al., 2021b). At all gestational ages, a very substantial layer of trabecular muscle was found in both ventricles (Fig. 10B). We invariably found that older specimens had a greater volume of trabecular muscle in each ventricle (Fig. 10C). The same findings were made for compact muscle, only the compact muscle increased in volume at a greater rate than the trabecular muscle (Fig. 10C). This difference in growth rate led to a decrease of the percentage of trabecular muscle (Fig. 10D), even if the absolute volume of trabecular muscle increased over time. Such proportional change has been construed as ‘compaction’ but given the positive growth of the trabecular layer there is no quantitative support for the addition of trabeculations to the compact wall. In adult shrew, trabeculations had a much greater volume than in any developmental stage and the right ventricle had more trabeculation than the left ventricle (3.90 mm^3^ and 1.32 mm^3^ respectively).

**Figure 10.**
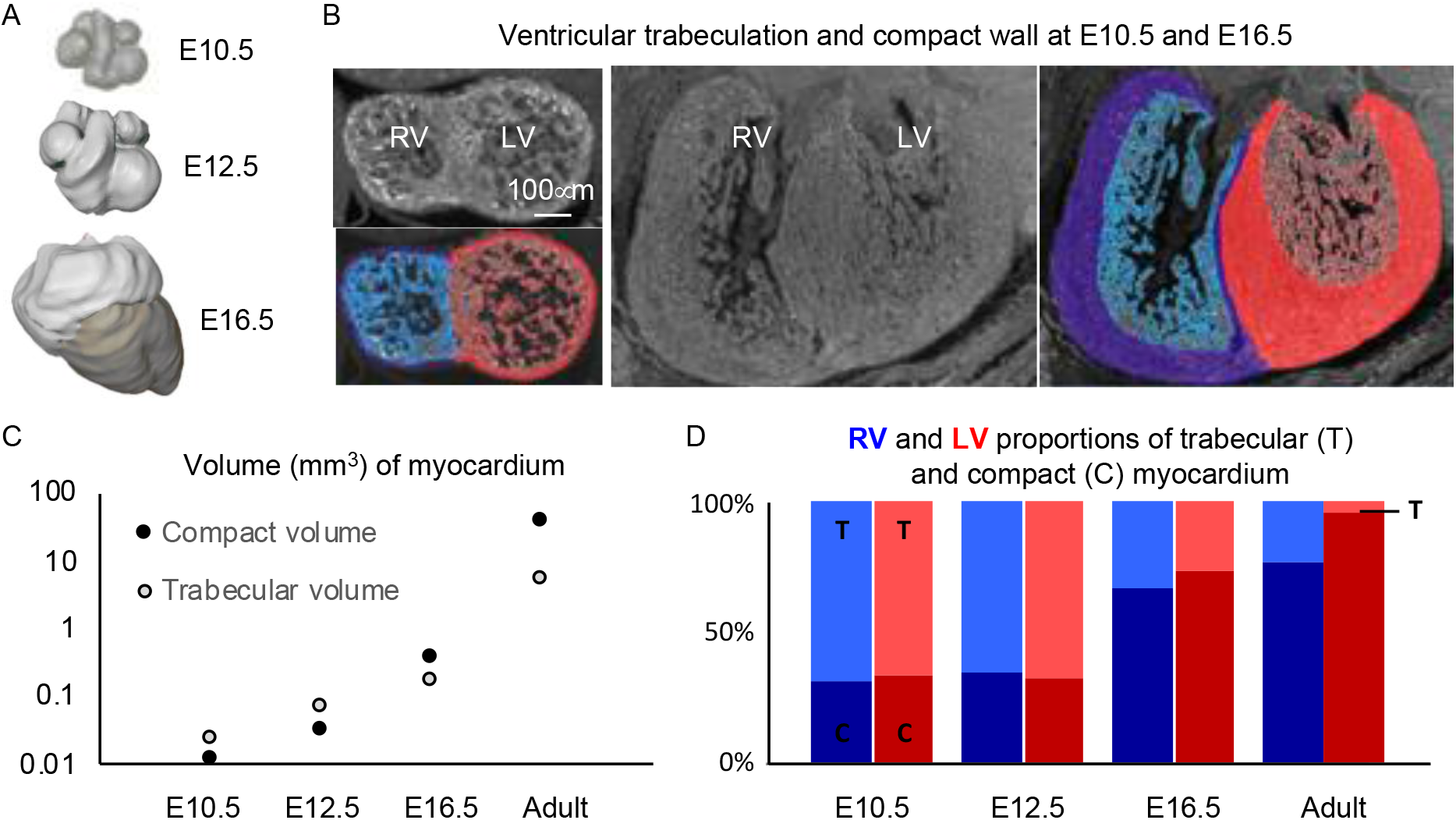
Development of ventricular trabeculation. **A**. Reconstruction based on micro-CT of a heart from each investigated embryonic age. **B**. Examples of histological sections and the subsequent labelling of these into right ventricular compact (blue) and trabecular layer (light blue) and left ventricular compact (red) and trabecular layer (light red). **C**. For each gestational age, the trabecular tissue volume increased for both ventricles, but not as fast as that of the compact tissue volume. **D**. The proportion of trabecular myocardium diminished with age and the ventricles thereby became less trabeculated even if there was no decrement in absolute volume of trabeculations.

#### The cardiac conduction system

Given the extremely high heart rates of shrews, it could be presumed that their hearts would contain a structurally pronounced conduction system. To investigate whether this was the case, we surveyed three histological series (Fig. 11A-B). A sinus node was found in the parietal and ventral junction of the right cranial caval vein and the right atrium (Fig. 11C). It was identifiable by being node-like in appearance, by being relatively rich in collagen and by a low expression of cardiac troponin I (Fig. 11D). In addition, Hcn4 which is a key marker of the conduction system (Boyett et al., 2021), was expressed in a subset of the myocardium with low expression of troponin I (Fig. 11D) which is another characteristic of conduction tissue (Sizarov et al., 2011). This presumed sinus node was small, approximately 100μm wide and 500μm dorso-ventrally long in all three investigated specimens. Concerning the atrioventricular conduction axis, an insulating plane disrupted the atrioventricular myocardial continuity in the right and left atrioventricular junction (Fig. 11E-F). At the base of the atrial septum, there was myocardium which was relatively dispersed by collagen and which expressed Hcn4 and expressed relatively little troponin I (Fig. 11G). This myocardium therefore at the appearance of the penetrating bundle of His. Ventrally, this myocardium came into continuity with the myocardium of the ventricular septal crest (Fig. 11H). We did not identify structures that resembled the atrioventricular node or bundle branches, but many of the sections that could have contained these structures also had artefacts from blood and connective tissue that had come loose during the staining procedures.

**Figure 11.**
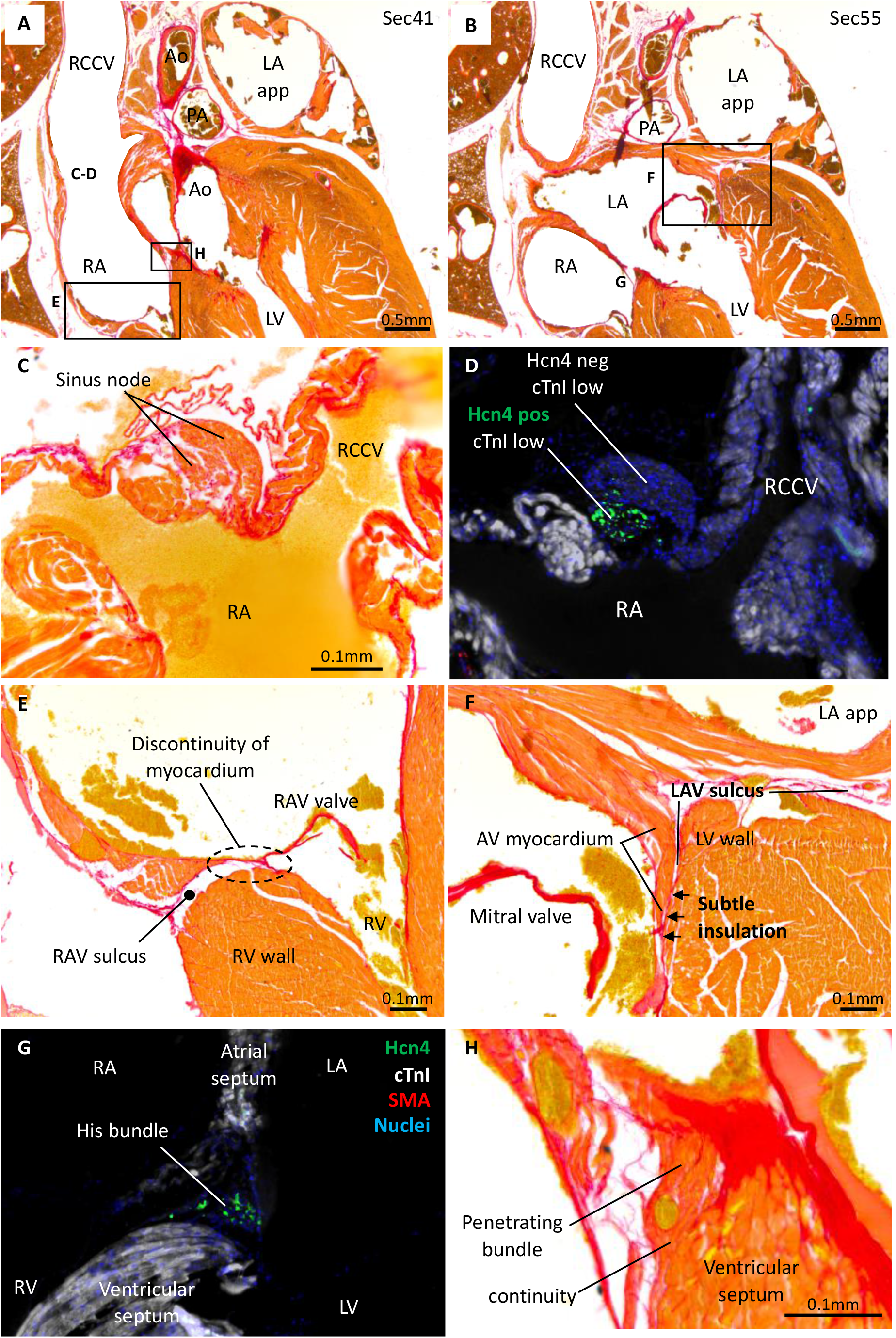
Cardiac conduction system. **A**. Overview of the sinuatrial (C-D) and right atrioventricular junction (E). **B**. Overview of the left atrioventricular junction (F), 280μm dorsal to image A. **C-D**. Sinus node on the junction of the right cranial caval vein (RCCV) and right atrium (RA), characterized by greater collagen infiltration (C) and expression of Hcn4 while having weak expression of cTnI (D). **E**. The right atrioventricular junction, showing no myocardial connection between the atrium and ventricle (RV) despite the junction contains very little tissue. **F**. The left atrioventricular junction, showing only a subtle amount of insulation between the atrioventricular (AV) and ventricular (LV) myocardium. **G**. Immunohistochemistry of the crest of the ventricular septum showing the His bundle (Hcn4 expression, weak cTnI expression) localized in a position much like the one indicated with ‘G’ in image B. **H**. The His bundle penetrates its insulation and becomes continuous with the ventricular septum. This image is a zoom-in of the region indicated with ‘H’ in image A. Ao, aorta; LA app, left atrial appendage; L(R)AV, left (right) atrioventricular; PA, pulmonary artery.

## DISCUSSION

The shrew heart is in most ways a typical mammal heart, despite being at the extreme lower end of the seven orders of magnitude that mammals range in size. Its exceptional traits can be summarized as its large relative size and great elongation, which has been reported before (Vornanen, 1989), and, as we show here, the extreme extent of the pulmonary venous myocardium. Sleeves of myocardium around the pulmonary veins are found in many mammals including human (Rowlatt, 1990). The sleeves are typically confined to the parts that are most proximal to the left atrium and they are thought to act as throttle valves of pulmonary venous return (Nathan and Gloobe, 1970). In the shrew, however, there is virtually no part of the pulmonary venous tree that is without a myocardial sleeve. It is therefore difficult to envision the role of throttle valve to this myocardium. In human, ectopic pacing of the atria often originates from the pulmonary venous myocardium and this setting requires pulmonary vein isolation by catheter-ablation. While electrocardiograms have been reported for shrews (Morrison et al., 1959, Nagel, 1986, Vornanen, 1989, Jurgens et al., 1996), it is not known whether their extensive pulmonary venous myocardium associates with a propensity to develop atrial arrhythmias.

### Traits that shrews share with most mammals

Shrews and most mammals have three caval veins, whereas the human setting of having a regressed left caval vein is less common (Rowlatt, 1990, Jensen et al., 2014a, Carmona et al., 2018). Compared to human, the caval vein myocardium is extensive, but many mammals have similarly extensive myocardium i.e. it extends to the vicinity of the pericardial border (Nathan and Gloobe, 1970, Jensen et al., 2014a). The number of pulmonary veins that connect to the left atrium is more variable in mammals than in other tetrapods (Kroneman et al., 2019) and two veins, as in shrews, is not uncommon (Rowlatt, 1990). The atrioventricular valves and tricuspid pulmonary and aortic valves of the shrews were typically of mammals (Rowlatt, 1990). In monotreme and marsupial mammals the right atrioventricular valve can be dominated by the parietal leaflet (Lankester, 1882, Runciman et al., 1992), but in the shrews the septal leaflet was well-developed as is common in eutherian mammals (Rowlatt, 1990). Eutherians are distinct from other mammals by having a second atrial septum which leaves a circular depression called the oval fossa on the right face of the atrial septum (Röse, 1890, Rowlatt, 1990, Jensen, 2019). The shrews also have an oval fossa and its dorso-cranial rim is provided by a fold in the atrial roof in a manner that much resembles the human setting (Anderson et al., 2014).

Given the extremely high heart rates of shrews, one could presume an unusual cardiac conduction system. The cardiac conduction has been identified on the basis of histological characters such as weak stain and richness in collagen (Davies, 1942, Ho et al., 2003) and on the presence of the funny current channel, Hcn4 (Sizarov et al., 2011, Boyett et al., 2021). We show in the shrew the presence of myocardium that is insulated by collagen and that expresses Hcn4 where a mammal sinus node and His bundle would be expected. A lot of our histology was perturbed by loose and displaced connective tissue and blood, and this hampered the identification of the atrioventricular conduction axis in particular. We therefore suggest that the absence of a clear identification of an atrioventricular node and bundle branches in our data should not be seen as strong evidence for the absence of these structures. From the data we do have, we consider it unlikely that the shrew cardiac conduction system is excessively developed.

### Unusual traits of the shrew heart

A right ventricular wall and septum almost free of trabeculations is not typical of mammals but it is found in shrews, as we report here, and bats, squirrels, and mustelids (Rowlatt, 1990). It is invariable so that the ventricles of embryos are highly trabeculated (Jensen et al., 2016), which has also been documented in shrews (Yasui, 1993). So a proportional change must take place from the embryonic and highly trabeculated setting to the adult setting which is virtually free of trabeculations. Two different processes have been proposed to explain such proportional change. One is that the trabecular and compact layers grow throughout development, but the growth rate of the two layers may differ periodically which then changes the layer proportions (Faber et al., 2021a). In agreement with this view, our data show the compact layer has a greater rate of growth than the trabecular layer and this reduces the proportion of trabecular muscle during gestation. The other proposed process is compaction, whereby trabeculations are removed from the trabecular layer and added to the compact wall (Rychterova, 1971, Chin et al., 1990). No decrement of the trabecular layer thickness has been documented however (Faber et al., 2021b), even though this is a predicted outcome if compaction takes place. In this light, the shrew ventricles are an interesting test for the hypothesis of compaction, because their right ventricular wall is comparative very smooth. The developmental data, however, do not support a role of compaction but it do support a role of differential growth rates.

### Unusual traits that may be ascribed to small size

One unusual trait of the shrew hearts was the little amount of fibro-fatty tissue that comprised the insulating plane between the atria and ventricles. In the left atrioventricular junction, the insulation was so meagre that it was difficult to assess whether the atrial and ventricular myocardium was in fact insulated from each other. In larger animals, millimeters of connective tissue separate the atrial and ventricular myocardium (Ho et al., 2003). Another unusual trait was the extremely large size of the main branches of the coronary vessels, when these were seen in proportion to the diameter of the aorta and thickness of the ventricular wall. In absolute size, the diameter of the coronary vessels was of course small and it was within the range of arterioles. In large conduit vessels such as the human aorta, diameter has almost no impact on resistance to blood flow as given by the Hagen-Poiseuille equation, whereas in the size range of arterioles, diameter change has a very large impact on resistance. In this light, the proportions of aorta-to-coronary-arteries found in human, for example, may not scale to the small size of shrews, because it would reduce the diameter of the coronary arteries so much as to render them high-resistance vessels.

### Highly unusual traits of the shrew heart

A very elongate ventricle is a highly unusual trait in a mammal and it is found in shrews (Vornanen, 1989) and the closely related moles of the genus *Talpa* (Rowlatt, 1968). Otherwise, elongate ventricles appear restricted to animals that are highly elongate themselves, such as snakes and caecilian amphibians (Ramaswami, 1944, Jensen et al., 2014b, de Bakker et al., 2015). Shrews and moles are not particularly elongate mammals and the significance of their highly unusual heart shape is not clear. The developing hearts that were investigated here, and previously (Yasui, 1993), were not much different in shape from developing mouse hearts and this suggests that the elongate shape develops late in gestation or even after birth.

Perhaps the most unusual morphological trait of the shrew heart is the extent of the pulmonary venous myocardium. It was previously demonstrated that pulmonary venous myocardium is present in shrews (Endo et al., 1997) and mice, for example, have myocardial sleeves that extend three bifurcations up the pulmonary venous tree (Mommersteeg et al., 2007). To the best of our knowledge, however, our report is the first documentation of the extreme extent of the shrew pulmonary venous myocardium which much exceeds that of mouse. The extreme extent aside, the amount of muscle around the pulmonary veins is not much, it is just a few percent of the total cardiac mass. Therefore, even if cardiac muscle is energetically very demanding (Mootha et al., 1997), the pulmonary venous myocardium may not necessarily impose a large metabolic cost and its advantage to organismal performance may not have to be great.

Observations on valves in veins have a long history (Franklin, 1927), but to the best of our knowledge this is the first report of a valve on the myocardial-venous boundary in the caudal caval vein. Whales have a sphincter at the same position (Lillie et al., 2018), but not a valve of leaflets. Even in reptiles, where the caval vein myocardium functions as a chamber and a valve in the caudal caval vein would seem advantageous (Jensen et al., 2017, Joyce et al., 2020), there is no valve. Because such valve is rarely sought after and it may easily be obscured by coagulated blood or by its collapse against the vessel wall, a dedicated investigation may be required to establish whether a valve in the caudal caval vein is highly unusual among vertebrates.

## Conclusion

In this study we investigated whether typically traits of mammal hearts scale to the size of shrews and for key anatomical traits, such as valves and septums, they do scale to the extremely small. Traits that do not scale may be the proportionally very large coronary arteries and the atrioventricular junction insulation, which comprise very little tissue. The pronounced elongation of the ventricle, the extreme extent of the pulmonary venous myocardial sleeves, and the valve in the caudal caval vein may set shrew hearts aside from other mammal hearts.

## ACKNOWLEDGMENTS

The authors would like to thank for her assistance Antonina Yu. Alexandrova with the collection of shrews, Quinn Gunst and Corrie de Gier-de Vries for their assistance with histology and immunohistochemistry and Jaco Hagoort for his assistance with Amira. The authors have no conflict of interest to declare.

## AUTHOR CONTRIBUTION

Conceptualization of the study was by BJ; acquisition of data and critical revision of the manuscript was by was by YHC, BIS, BJ; data analysis and interpretation, and drafting of the manuscript was by YHC and BJ.

## DATA AVAILABILITY STATEMENT

The data that support the findings of this study are available from the corresponding author upon reasonable request.

